# Molybdenum-based diazotrophy in a *Sphagnum* peatland in northern Minnesota

**DOI:** 10.1101/114918

**Authors:** Melissa J. Warren, Xueju Lin, John C. Gaby, Cecilia B. Kretz, Max Kolton, Peter L. Morton, Jennifer Pett-Ridge, David J. Weston, Christopher W. Schadt, Joel E. Kostka, Jennifer B. Glass

## Abstract

Microbial N_2_ fixation (diazotrophy) represents an important nitrogen source to oligotrophic peatland ecosystems, which are important sinks for atmospheric CO_2_ and susceptible to changing climate. The objectives of this study were: (i) to determine the active microbial group and type of nitrogenase mediating diazotrophy in a ombrotrophic *Sphagnum*-dominated peat bog (the S1 peat bog, Marcell Experimental Forest, Minnesota, USA); and (ii) to determine the effect of environmental parameters (light, O_2_, CO_2_, CH_4_) on potential rates of diazotrophy measured by acetylene (C_2_H_2_) reduction and ^15^N_2_ incorporation. Molecular analysis of metabolically active microbial communities suggested that diazotrophy in surface peat was primarily mediated by *Alphaproteobacteria* (*Bradyrhizobiaceae* and *Beijerinckiaceae*). Despite higher dissolved vanadium (V; 11 nM) than molybdenum (Mo; 3 nM) in surface peat, a combination of metagenomic, amplicon sequencing and activity measurements indicated that Mo-containing nitrogenases dominate over the V-containing form. Acetylene reduction was only detected in surface peat exposed to light, with the highest rates observed in peat collected from hollows with the highest water content. Incorporation of^15^N_2_ was suppressed 90% by O_2_ and 55% by C_2_H_2_, and was unaffected by CH_4_ and CO_2_ amendments. These results suggest that peatland diazotrophy is mediated by a combination of C_2_H_2_ - sensitive and C_2_H_2_ - insensitive microbes that are more active at low O_2_ and show similar activity at high and low CH_4_.

**Importance:** Previous studies indicate that diazotrophy provides an important nitrogen source and is linked to methanotrophy in *Sphagnum*-dominated peatlands. However, the environmental controls and enzymatic pathways of peatland diazotrophy, as well as the metabolically active microbial populations that catalyze this process remain in question. Our findings indicate that oxygen levels and photosynthetic activity override low nutrient availability in limiting diazotrophy, and that members of the *Alphaproteobacteria* (*Rhizobiales*) catalyze this process at the bog surface using the molybdenum - based form of the nitrogenase enzyme.

## Introduction

High-latitude peatlands store approximately one-third of global soil carbon and may pose a climatic threat if rising global temperatures accelerate the release of this stored carbon in gaseous forms, as either carbon dioxide or methane (35, 64, 106). Mineral-poor (ombrotrophic) peatlands receive most of their nutrient inputs from atmospheric deposition and contain *Sphagnum* moss as their primary plant cover (12, 64). The peatmoss *Sphagnum* is a keystone genus in these ecosystems and is responsible for much of the primary production and recalcitrant dead organic matter (13, 103). *Sphagnum* mosses also host complex microbiomes (9, 58, 78, 79), including N_2_-fixers (diazotrophs) that are significant nitrogen sources to peatland ecosystems (5).

Despite decades of research, there is still much debate on the identity of the dominant diazotrophs in ombrotrophic peatlands. Early work implicated *Cyanobacteria* (2, 36, 37) or heterotrophic bacteria (88) based primarily on microscopic studies, while more recent molecular analyses argue for the importance of methanotrophic *Beijerinckiaceae* (19) as major diazotrophs in *Sphagnum* peat bogs (8, 22, 44, 99). Possible contributions from other potential diazotrophs, such as strictly anaerobic methanogenic *Euryarchaeota*, remain unknown. However, it is quite possible that diverse diazotrophs exist within defined niches of peatland environments (60).

Diazotrophy is catalyzed by the nitrogenase metalloenzyme, a complex of thee subunits (H, D and K) that contains abundant iron as Fe-S clusters. This enzyme is extremely O_2_ sensitive (105), and must be protected from exposure to O_2_ for diazotrophy to occur (25). The most common form of nitrogenase, encoded by *nif* genes, contains molybdenum (Mo) as its cofactor. When Mo is scarce, some species of Bacteria and Archaea express nitrogenases containing vanadium (V; *vnf* genes) or iron (Fe; *anf* genes) in place of Mo, but these “alternative” nitrogenases are less efficient than the Mo form (74, 101). The most conserved nitrogenase gene, *nifH* (77), has become the marker gene of choice for environmental diazotrophy (28, 29, 107). Phylogenetic studies show five *nifH* clusters: aerobic bacteria (cluster I); alternative nitrogenases (cluster II); anaerobic bacteria and archaea (cluster III); uncharacterized sequences (cluster IV), and paralogs related to chlorophyll biosynthesis (cluster V; 85). Because *vnfH* and *anfH* genes in cluster II cannot be differentiated by sequence alone, the D-subunit (*nifD* / *vnfD* / *anfD*) has become the preferred marker gene for studies of alternative nitrogenases (4). Consistent with higher concentrations of V than Mo in most rocks (102), microbes from diverse soils have been shown to contain *vnfD* genes (4, 7, 18, 45, 70, 73). Given that oligotrophic conditions dominate in peatlands, trace metals may limit diazotrophy. However, little is known about trace metal availability and the role of alternative nitrogenase pathways in ombrotrophic peatlands.

Similarly, methane monooxygenase (MMO, the enzyme that catalyzes the first step of methane oxidation) occurs in particulate (copper (Cu)-containing pMMO) and soluble (Fe-containing sMMO) forms. While pMMO has more specific substrate requirements, pathways that employ sMMO can use a wider range of compounds (15). Both forms of MMO are inhibited by acetylene (C_2_H_2_) (14, 82). In organisms with both sets of genes, pMMO is expressed when Cu is abundant, whereas Cu limitation induces sMMO expression (90). The dominant peatland methanotrophs in *Alphaproteobacteria* and *Gammaproteobacteria* tend to possess both MMOs (11, 24, 39, 52, 68), although *Methylocella* and *Methyloferula* species containing solely sMMO have been isolated from peat bogs (20, 21, 100). While most studies have primarily targeted the *pmoA* gene (24, 52), *mmoX* genes and transcripts have also been reported in peatlands (63, 68, 84), raising questions about the relative importance of each form for peatland methane oxidation.

The acetylene reduction assay (ARA) is commonly used as a proxy for diazotroph activity (41, 42). This assay is effective for capturing the potential activity of diazotrophs that are not inhibited by C_2_H_2_, such as *Cyanobacteria* and non-methanotrophic *Proteobacteria* (e.g. *Bradyrhizobiaceae*) (50). However, a number of functional guilds of microorganisms, including methanotrophs, methanogens, sulfate reducers, and nitrifiers, are inhibited by C_2_H_2_ (16, 49, 80, 81, 92, 94. If these or other C_2_H_2_-sensitive microbes perform diazotrophy and/or provide substrates to other diazotrophs (see Fulweiler, et al. (27)), ARA may underestimate diazotrophy in that system. Thus, recent studies have shifted to tracking diazotrophy by incorporation of the stable isotope tracer, ^15^N_2_ (55, 60, 61, 99).

In this study of the S1 peat bog at the Marcell Experimental Forest in northern Minnesota, USA, dissolved macro-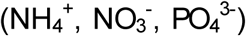 and micro-(Fe, Cu, V, Mo) nutrients were profiled along with the community composition and abundance of diazotrophic microorganisms. We also performed separate laboratory incubation experiments to measure potential rates of ARA and ^15^N_2_ incorporation in order to: (a) assess environmental controls (light, O_2_, CH_4_) on diazotrophy; (b) quantify the effect of C_2_H_2_ on rates of diazotrophy and methanotrophy; and (c) search for diagnostic markers for alternative nitrogenase activity such as a low conversion factor of ARA to ^15^N_2_-incorporation (4) and C_2_H_2_ reduction to ethane (23). Finally, we make recommendations on universal *nifH* primers for amplicon sequencing and quantitative PCR based on our findings.

## Materials and Methods

### Site description and sample collection

Samples were collected from the S1 (black spruce-*Sphagnum* spp.) peat bog at Marcell Experimental Forest (MEF; 47°30.476’ N; 93°27.162’ W), the site of the DOE SPRUCE (Spruce and Peatland Responses Under Climatic and Environmental Change) experiment in northern Minnesota, USA (40). The S1 bog is ombrotrophic and acidic (average pH 3.5-4; 57, 89). Over the summer months, the water table is ±5 cm from the hollow surface (38, 89). Dissolved O 2 levels decrease to below detection (∽20 ppb) within the top 5 cm of the bog. Three locations were sampled along S1 bog transect 3 (T3) at near, middle and far sites (see Lin, et al. (69) for further details). Surface (0-10 cm depth) peat was collected from hollows dominated by a mixture of *Sphagnum fallax* and *S. angustifolium*, and hummocks dominated by *S. magellanicum*. Peat depth cores (0-200 cm) were sampled from hollows where the water level reached the surface of the *Sphagnum* layer.

### Macronutrients

Peat porewater was collected using piezometers from 0, 10, 25, 50, 75, 150 and 200 cm depth. Piezometers were recharged the same day as collection, and porewater was pumped to the surface, filtered through sterile 0.2 μm polyethersulfone membrane filters, and stored frozen until analysis. Nitrate 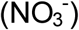 and nitrite 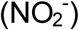 were analyzed using the spectrophotometric assay as described by García-Robledo, et al. (31). Ammonium 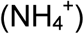 concentrations were determined with the indophenol blue assay (95). Phosphate concentrations were measured with the molybdate-antimony ascorbic acid colorimetric assay (75).

### Micronutrients

Peat porewater was collected from two locations in the S1 bog from cores at 0-30 cm, 30-50 cm, and 100-150 cm depths by filtration through 0.15 μm Rhizon soil samplers (Rhizosphere Research Products). All plastics were washed with HCl prior to sampling; Rhizon soil samplers were cleaned by pumping 10 mL of 1 N HCl through them, followed by a rinsing with ultrapure water until the pH returned to neutral (∽100 mL / filter). After collection, samples were acidified to 0.32 M HNO_3_ (Fisher Optima) and analyzed using a Thermo ELEMENT 2 HR-ICP-MS (National High Magnetic Field Laboratory, Florida State University). Initial analyses resulted in frequent clogging of the nebulizer, likely due to abundant dissolved organic carbon. Therefore, samples were diluted 1:10 to minimize interruptions from nebulizer clogs. Concentrations were quantified with a 7-point external calibration using standards prepared in 0.32 N HNO_3_ from a multi-element standard mix (High Purity Standards).

In order to generate an organic-free sample matrix suitable for ICP-MS analysis without contaminating or diluting the sample, subsequent samples were digested as follows: 1 mL aliquots of the porewater samples were heated in 15-mL Teflon beakers (Savillex) with 1 mL of 16 N HNO_3_ (Ultrex II, JT Baker) and 100 μL of 30% H_2_O_2_ (Ultrex II, JT Baker) for 36 h at 230°C in a trace metal clean, polypropylene exhaust hood. The HNO_3_/H_2_O_2_ mixture oxidizes any DOM to CO_2_, but the resulting matrix is too acidic for direct ICP-MS introduction. Therefore, samples were evaporated to near dryness and resuspended in a 0.32 N HNO_3_ matrix suitable for ICP-MS analysis, and analyzed by ELEMENT2 ICP-MS along with parallel blank solutions.

### Quantification and sequencing of gene and transcript amplicons

Peat was frozen on dry ice at the field site in July 2013, or in liquid N_2_ after 7 day incubations at 25°C in the light under degassed (80% N_2_ + 20% CO_2_) headspace with 1% C_2_H_2_, with or without 1% CH_4_ for June 2014 incubations (see ARA section). DNA and RNA were extracted with MoBio PowerSoil DNA and total RNA extraction kits, respectively, as described in Lin, et al. (68). RNA was cleaned with a TURBO DNA-free kit (Ambion). Nucleic acid purity was analyzed for 260/280 absorbance ratio (1.8-2.0) on a NanoDrop spectrophotometer. cDNA was synthesized using the GoScript reverse transcription system (Promega) according to the manufacturer’s protocol.

Plasmid standards for qPCR were constructed according to Lin, et al. (66). Primer pairs are given in Table 1. The gene fragments of *nifH*, *pmoA* and *mcrA* for constructing plasmid standards for qPCR were amplified from genomic DNA of *Rhodobacter sphaeroides*, *Methylococcus capsulatus* Bath, and S1 peat bog peat soil, respectively. To prepare cDNA standards, plasmid DNA with a positive gene insert was linearized with NcoI restriction enzyme following the manufacturer’s protocol (Promega), and purified by MinElute PCR purification kit (Qiagen). RNA was synthesized from the linearized plasmid DNA using the Riboprobe *in vitro* transcription system according to the manufacturer’s protocol (Promega), followed by cDNA synthesis using the GoScript reverse transcription system (Promega) according to the manufacturer’s protocol.

**Table 1.**
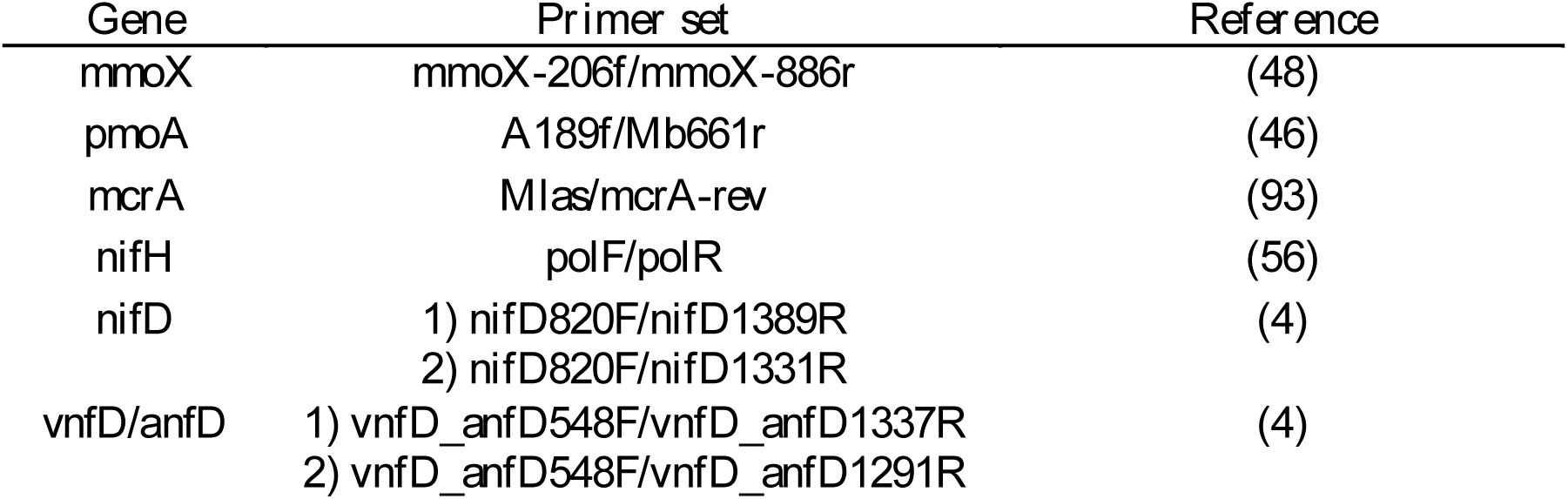
qPCR and sequencing primers used in this study. PCR conditions were based on refs. (24), (63), (93), (29), and (4) for *pmoA, mmoX, mcrA*, *nifH*, and *nifD / vnfD / anfD*, respectively.

| Gene | Primer set | Reference |
| --- | --- | --- |
| *mmoX* | mmoX-206f/mmoX-886r | (48) |
| *pmoA* | A189f/Mb661r | (46) |
| *mcrA* | Mlas/mcrA-rev | (93) |
| *nifH* | polF/polR | (56) |
| *nifD* | 1) nifD820F/nifD1389R<br>2) nifD820F/nifD1331R | (4) |
| *vnfD/anfD* | 1) vnfD_anfD548F/vnfD_anfD1337R<br>2) vnfD_anfD548F/vnfD_anfD1291R | (4) |

The abundance of functional gene transcripts were quantified in samples run in duplicate on a StepOnePlus Real-Time PCR System (ABI) using Power SYBR Green PCR master mix. Reaction mixtures were 20-μL reaction mixtures with 2-μL of template cDNA (10-100ng/μL) added to 10 μL of SYBR green master mix, 0.5-1.6 μL of each forward and reverse primer (0.3-0.8 μM final concentration; Table 1), and 4.8-6.5 μL of PCR-grade water. Samples were run against a cDNA standard curve (10^1^ to 10^7^ copies of plasmid gene fragment) on a StepOnePlus qPCR instrument with 96 wells with an initial denaturation step of 2-5 min at 95°C and 40 cycles of denaturation at 95°C for 15-30 s, annealing at 55-64°C for 30-45 s, extension at 72°C for 30-45 s, and data acquisition at 83-86.5°C for 16-30 s. To minimize the effects of inhibitors in assays, peat DNA was diluted to 1/40 of original concentrations, and duplicate 20-μL reaction mixtures, each containing 2-μL of diluted DNA, were run for each sample. Functional gene and transcript copy numbers were normalized to dry weight of peat, or 16S rRNA transcript copies for incubation samples. Amplicons were sent to the University of Illinois at Chicago for DNA sequencing using a 454 platform. Raw sequences were demultiplexed, trimmed, and quality filtered in CLCbio. The phylogeny of *vnfD/anfD* sequences was inferred using the Maximum Likelihood method based on the Kimura 2-parameter model in MEGA5 (96).

### Acetylene reduction and methane consumption rates

Samples of bulk peat (*Sphagnum* spp. and surrounding soil) were collected from 0-10 and 10-30 cm depths in September 2013, April 2014, June 2014, September 2014 and August 2015, and stored at 4**°** C until the start of laboratory incubations. Samples from 0-10 cm depth were gently homogenized so as not to rupture *Sphagnum* spp. tissues, while peat samples from 10-30 cm depth were fully homogenized. For each sample, 5 g of bulk peat were placed in 70 mL glass serum bottles, stoppered with black bromobutyl stoppers (Geo-Microbial Technologies, pre-treated by boiling 3x in 0.1 M NaOH) and sealed with an aluminum crimp seal. Headspaces were oxic (room air, 80% N_2_ + 20% O_2_) or degassed (100% N_2_ or 80% N_2_ + 20% CO_2_) with or without 1% C_2_H_2_ or 1% CH_4_. Treatments were incubated for one week at 25°C in the light or dark. A gas chromatograph with a flame ionization detector (SRI Instruments) equipped with a HayeSep N column was used to quantify CH_4_, C_2_H_2_ and C_2_H_4_. Samples were measured for C_2_H_4_ production daily until C_2_H_4_ production was linear (∽7 days). Controls not amended with C_2_H_2_ did not produce ethylene (C_2_H_4_). Incubations of hollow peat from June 2014 incubated in oxic headspace with and without 1% C_2_H_2_ were also monitored for consumption of 1% CH_4_. Statistical analysis was performed with JMP Pro (v. 12.1.0) using the Tukey-Kramer HSD comparison of all means.

**^*15*^*N*_*2*_ *incorporation rates.*** In September 2014, samples were quantified for N_2_ fixation rates by ^15^N_2_ incorporation in parallel with ARA measurements. Incubations were set up as described above and supplemented with 7 mL of 98% ^15^N_2_ (Cambridge Isotope Laboratories, Tewksbury, MA, USA). After 7 days, samples were dried at 80°C, homogenized into a fine powder, and analyzed for N content and δ^15^N by isotope-ratio mass spectrometry (IRMS) with a MICRO cube elemental analyzer and IsoPrime100 IRMS (Elementar) at the University of California, Berkeley, corrected relative to National Institute of Standards and Technology (NIST, Gaithersburg, MD, USA) standards.

### Metagenomic analyses

Metagenomes were generated in a previous study (67). Diazotrophic and methanotrophic pathways were investigated using the following bioinformatics approaches. Briefly, Illumina reads were filtered by quality (Phred33 score threshold of Q25) using Trim Galore (Babraham Bioinformatics) and a minimum sequence length cut off of 100 bp. The sequences were then queried using RAPSearch2 (109) against the NCBI-nr database of non-redundant protein sequences as of November 2013. Sequences with a bit-score of 50 and higher were retained to determine the total number of functional genes for normalization across the different samples. The taxonomic composition of protein-coding sequences was determined based on the taxonomic annotation of each gene according to the NCBI-nr taxonomy in MEGAN5 (47) (min score: 50; max expected: 0.01; top percent: 10; min complexity: 0.3).

To classify sequences by nitrogenase cluster type, genes were analyzed using BLASTX (e-value 0.1; bit-score 50) versus a custom *nifH* database that includes a phylogenetic tree to distinguish the principal clusters (I, V, III) in the *nifH* phylogeny, as well as paralogous cluster IV, *nifH*-like sequences (28). *nifH* genes from the four clusters were normalized to total protein-coding genes from RAPSearch2 output sequences. The relative abundance of particulate (*pmoA*) vs. soluble (*mmoX*) methane monooxygenase was based on previous analyses in Lin, et al. (68).

## Nucleotide sequence accession numbers

Metagenomes were reported in a previous study (67) and deposited in BioProject PRJNA382698 (SAMN06712535-06712540). *pmoA* cDNA amplicons were reported in a previous study (24) and deposited in BioProject PRJNA311735. *nifH*, *mcrA*, *nifD*, and *vnfD/anfD* cDNA amplicons were deposited in BioProjects PRJNA382268, PRJNA382282, PRJNA382288 and PRJNA382295, respectively.

## Results

### Macro-and micro-nutrient profiles

In S1 bog hollows, 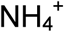 was ∽2 μM from the surface to 25 cm depth, and increased at greater depths (Fig. 1; Fig. S1a). Nitrate was <1 μM in surface peat and decreased with depth (Fig. S1b). Phosphate was <0.1 μM from the surface to 25 cm depth, and then increased with depth (Fig. S1c). For the metal pairs of greatest interest to this study, V (5-21 nM) was consistently more abundant than Mo (1-7 nM; Fig. 3), and Fe (7-35 μM) was three orders of magnitude higher than Cu (7-38 nM; Fig. S3) at all three depth intervals (0-30, 30-50 and 100-150 cm), and at three sampling dates (September 2014, June 2015, and September 2015; data shown for June 2015 in Figs. 3 and S2). Other trace nutrients were in the nano-to micro-molar range: Co (5-20 nM), Ni (10-80 nM), Zn (50-250 nM) and Mn (60-2220 nM; Table S1). Macro-and micro-nutrient profiles were essentially identical in profiles from Zim bog, another ombrotrophic bog ∽80 km southeast of the S1 bog, sampled in September 2014, with the exception of lower Fe at the surface (data not shown).

**Figure 1.**
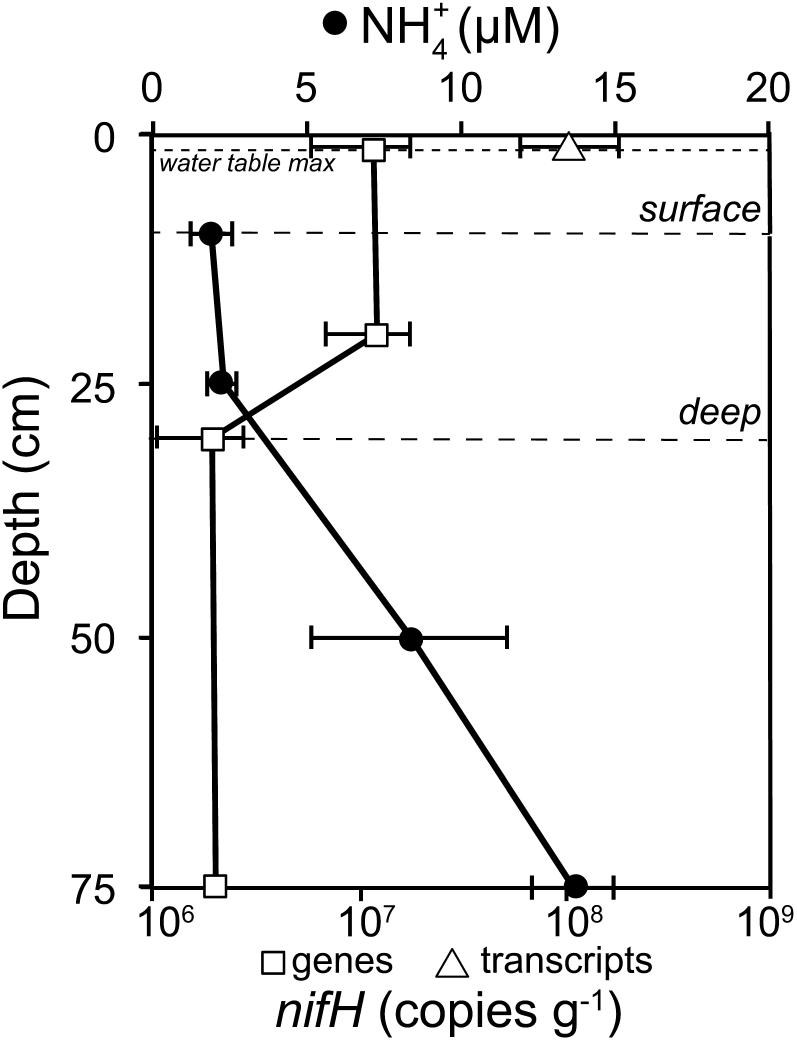
Depth profiles of NH_4_^+^ concentrations (black circles) and *nifH* gene copies (white squares) and transcripts (white triangle) in units of copies per gram of dry weight for the S1 bog T3 mid site. Ammonium concentrations are means of measurements from May, June and September 2014. *nifH* copy numbers are from July 2013. Error bars are standard errors. *nifH* transcripts were not detected at 20, 30 and 75 cm depths. Surface (0-10 cm) and deep (10-30 cm) peat depth intervals used for rate measurements are designated by dashed lines. The maximum water table depth at the S1 site in July 2013 was 2 cm below the hollow surface (dotted line) (38).

### Nitrogenase expression and phylogeny

With the polF/polR primer pair, we measured1.2x10^7^ copies g^−1^ *nifH* genes at 1 and 20 cm, and 0.2x10^7^ copies g^−1^ at 30 and 75 cm; *nifH* transcripts (12.2 x 10^7^ copies g^−1^) were only detected at 1 cm (10:1 transcript: gene ratio), and not at deeper depths (Fig. 1; Fig. S3a). Sequencing of cDNA from surface peat amplified with polF/polR (*nifH*) and nifD820F/nifD1331R (*nifD*) primers showed that the majority of nitrogenase transcripts belonged to cluster I (*Alphaproteobacteria*), with *Beijerinckiaceae* dominating *nifH* sequences and *Bradyrhizobiaceae* dominating *nifD* sequences (Fig. 2). Additional alphaproteobacterial *nif* transcripts matched to *Rhodospirillaceae, Rhizobiaceae, Rhodobacteraceae, Methylocystaceae* and *Xanthobacteraceae* (Fig. 2). *Gammaproteobacteria*, *Cyanobacteria* (*Oscillatoriophycideae*) and *Nitrospira* were also observed at lower abundance in cDNA amplicon libraries (data not shown).

**Figure 2.**
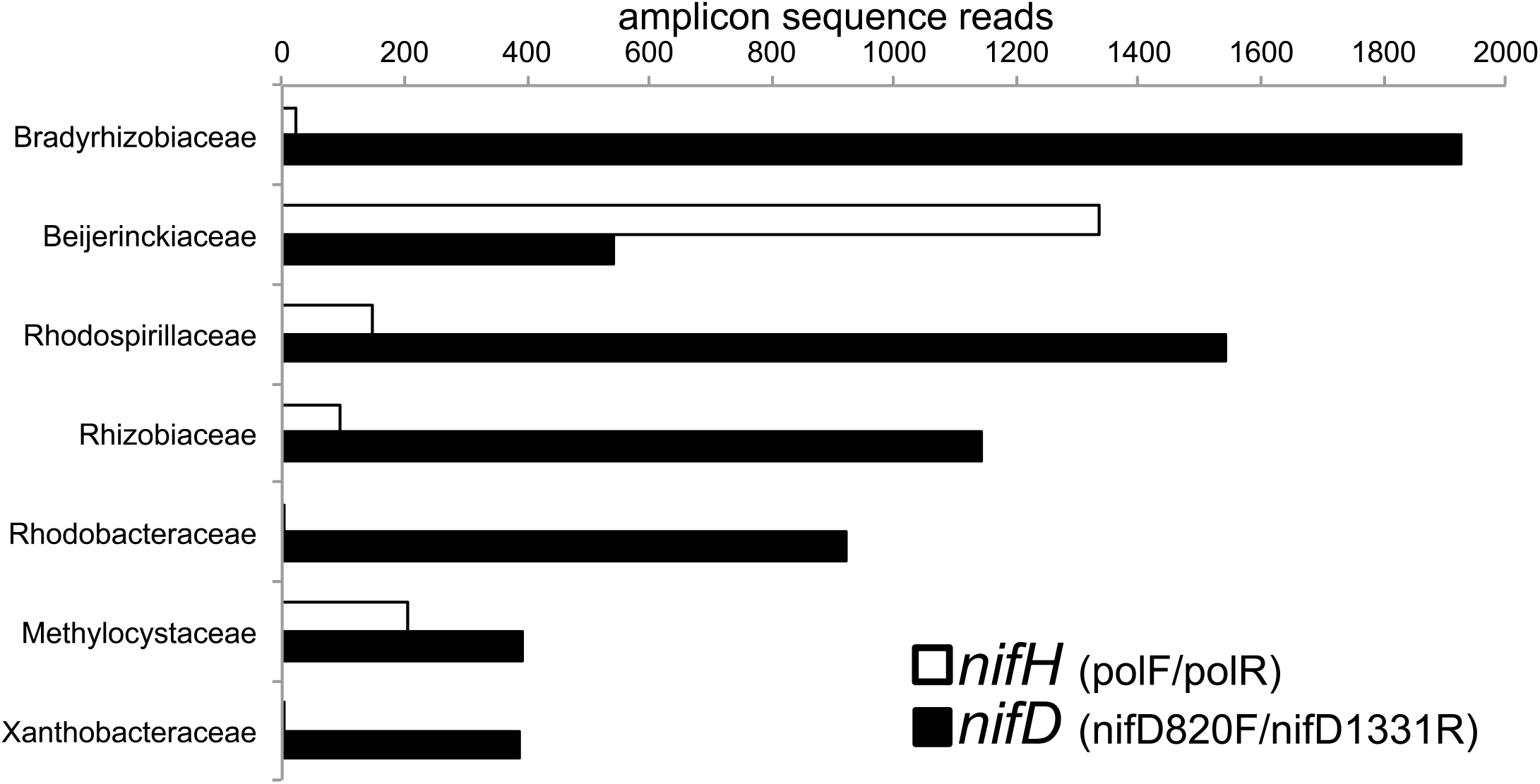
Numbers of cDNA amplicon sequence reads for *nifH* and *nifD* alphaproteobacterial transcripts. Primer sets were polF/polR and nifD820F/nifD1331R for *nifH* and *nifD*, respectively.

**Figure 3.**
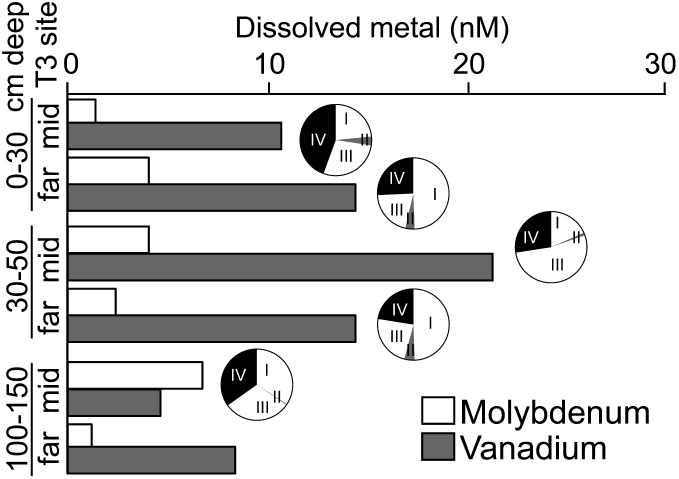
Dissolved molybdenum (white) and vanadium (gray) concentrations in pore water from three depths in S1 peat hollows (mid and far sites along T3 transect) from June 2015. Pie charts show the relative abundance of genes encoding the five nitrogenase H-subunit clusters from metagenomes for each depth; clusters I and III encode Mo-Fe nitrogenases (*nifH)*; cluster II encodes alternative (*vnfH, anfH*) nitrogenases; cluster IV encode nitrogenase paralogs. Deepest metagenomes were from 75 cm; insufficient numbers of nitrogenase H-subunit sequences were recovered from the far site for cluster analysis.

In metagenomes, *nifH* genes were roughly equally distributed between Mo-dependent clusters I and III, and cluster IV/V paralogs (Fig. 2; Table S2). Sequences from Cluster II (alternative nitrogenases, *vnfH* / *anfH*) were scarce at all depths (<5% overall); two *vnfD* genes from metagenomes showed phylogenetic similarity to those from soil *Proteobacteria* (Fig. S4). Attempts to amplify *vnfD/anfD* from cDNA yielded few reads; those recovered were most similar to *Alphaproteobacteria anfD* from *Rhodopseudomonas* species (Fig. S4).

### Methane-related gene expression and phylogeny

Like *nifH*, particulate methane monooxygenase (*pmoA)* and methyl coenzyme M reductase (*mcrA*) transcripts showed the highest abundance in surface peat (Fig. S3b,c). Surface *pmoA* transcripts mapped to *Methylocystaceae* (75%) and *Methylococcaceae* (25%). Surface *mcrA* transcripts mapped to *Methanosarcina* (58%), *Methanocella* (28%), and *Methanoregula* (11%). Attempts to amplify *mmoX* from cDNA were unsuccessful (data not shown). In metagenomes, genes for *pmoA* were dominant in surface peat, whereas the relative abundance of *mmoX* sequences increased with depth (Fig. S2).

### Rates of diazotrophy and methanotrophy

Potential rates of acetylene reduction were measured for peat collected from S1 bog hollows and hummocks in April, June, August and September 2013-2015 and incubated for 1 week at 25°C. Acetylene reduction to ethylene was only detected in surface (0-10 cm) peat samples incubated in the light, and not in deep (10-30 cm) peat, nor in surface peat incubated in the dark. Acetylene reduction to ethane was not detected in any incubation (data not shown). *Sphagnum* in peat incubations exposed to light became visibly greener over the course of the incubation (Fig. 4).

**Figure 4.**
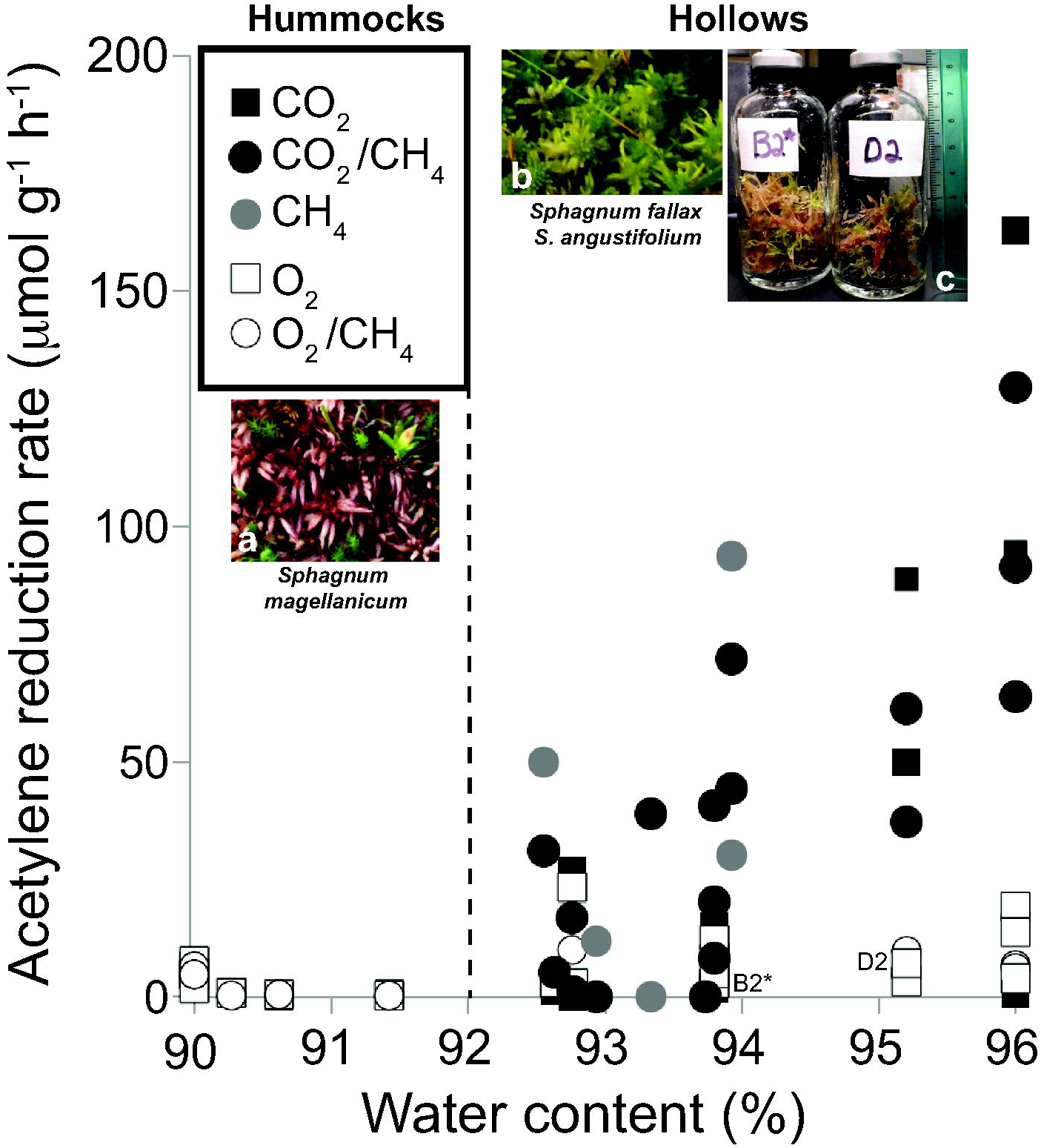
Acetylene reduction rates for hummocks (90-91% water content) and hollows (93-96% water content) at the S1 bog, T3 transect (0-10 cm depth incubated in the light at 25°C for 7 days). ARA units are μmol ethylene produced per gram of dry weight per hour. Photo insets show dominant *Sphagnum species* in (a) hummocks (*S. magellanicum*) and (b) hollows (*S. fallax* and *S. angustifolium*). Photo inset (c) shows *Sphagnum* greening after incubation of hollow samples in the light for 7 days at 25°C; bottle B2* (April 2014) received air headspace with 1% CH_4_ and treatment D2 (Sept 2013) received air headspace without CH_4_. The vertical dotted line divides the hummock samples (dominated by S. *magellanicum*) from the hollow samples (dominated by *S. fallax / angustifolium*) in terms of water content (92%).

In hollows, where surface peat was dominantly covered by a mix of *S. fallax* and *S. angustifolium*, ARA rates were higher and more variable in degassed vs. oxic incubations (0-163 vs. 2-23 μmol C_2_H_4_ g^−1^ h^−1^), and were unaffected by the presence or absence of 20% CO_2_. ARA in hollows incubated with degassed headspace was positively correlated (*P* < 0.0001) with peat water content (93 to 96%). In both hollows and hummocks, ARA rates were not affected by addition of 1% CH_4_. In hummocks with lower water content (90-91%), surface peat was dominantly covered by *S. magellanicum*, and oxic and degassed treatments had similarly low ARA rates (0-8 μmol C_2_H_4_ g^−1^ h^−1^). *nifH* transcripts in surface peat from hollows incubated under degassed headspace with 1% C_2_H_2_, with or without 1% CH_4_, ranged from 10^4^−10^7^ copies g^−1^ (July 2014; data not shown), which was 1-4 orders of magnitude lower than field samples from the previous summer (10^8^ copies g^−1^; Fig. 1), and higher and more variable in hollows than *nifH* transcripts from hummock incubations (Fig. S5).

^15^N_2_ incorporation showed similar overall trends as ARA (e.g. 90% higher rates in degassed vs. oxic conditions, and no significant CH_4_ effect; Fig. 5). In degassed treatments, 1% C_2_H_2_ inhibited ^15^N_2_ incorporation by 55% but had no effect on oxic treatments. In oxic treatments, CH_4_ consumption rates were 100 times higher than ^15^N_2_ incorporation rates, and 1% C_2_H_2_ addition suppressed CH_4_ oxidation rates by 95% (Fig. 5). Using the four sites measured with both methods, a conversion factor of 3.9 for ^15^N_2_-to-ARA was calculated (Fig. S6). In sum, laboratory incubations of native peats revealed that diazotrophy was stimulated by light, suppressed by O_2_, and minimally affected by CH_4_ and CO_2_.

**Figure 5.**
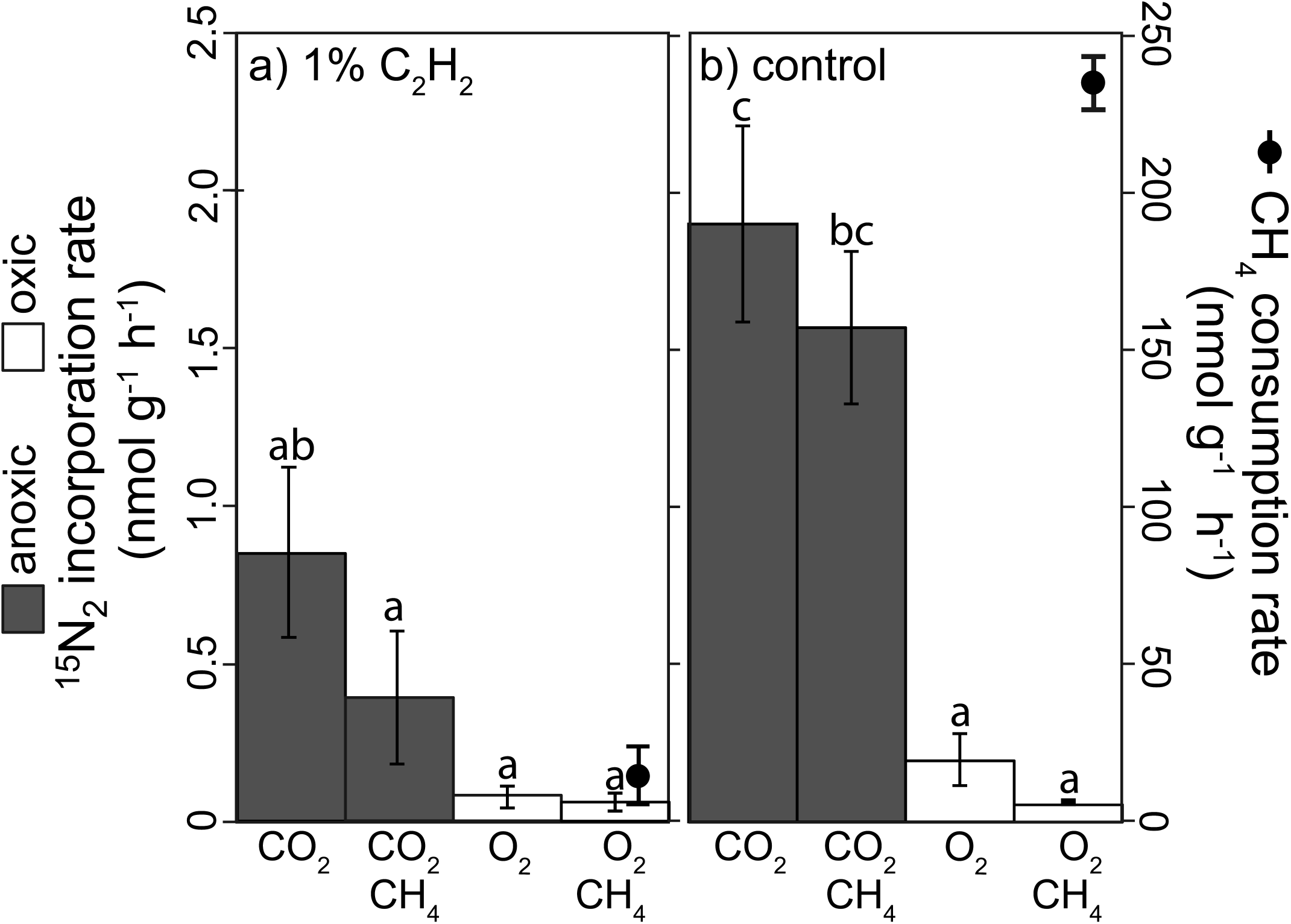
Effect of 1% C_2_H_2_ on ^15^N_2_ incorporation and CH_4_ consumption for S1 bog surface peat. Rates were measured for samples collected from the NW S1 bog transect in September 2014. ^15^N_2_ incorporation treatment conditions were 80%N_2_ + 20% CO_2_±1% CH 4 (shaded bars) or 80%N_2_ + 20% O_2_±1% CH 4 (white bars), with (a) and without (b) 1% C_2_H_2_; units are nmol ^15^N_2_ incorporated per gram of dry weight per hour. CH_4_ consumption treatments were 80%N_2_ + 20% CO_2_ + 1% CH_4_ (black circles) with (a) and without (b) 1% C_2_H_2_; units are nmol CH_4_ consumed per gram of dry weight per hour. Error bars are standard errors. Different letters indicate statistically different elemental contents (p < 0.05 based on Tukey-Kramer HSD test).

## Discussion

### Diazotrophs are active in surface peat

By definition, the only source of nutrients to ombrotrophic peat bogs is the atmosphere (65). Scarce (low μM) dissolved nitrogen and phosphorus in S1 peat pore water suggests oligotrophic conditions (65, 89, 97), consistent with low rates of atmospheric deposition in Minnesota peat bogs (43). Low nutrient concentrations add further evidence to previous suggestions that these nutrients may limit *Sphagnum* productivity (60) and/or complex mechanisms may exist for nutrient scavenging at ultra-low concentrations. Diazotroph activity (ARA and *nifH* transcription) was solely detected in surface peat samples incubated in the light. The apparent absence of diazotrophy at greater depths is consistent with previous reports (86), and may be due to light limitation and/or remineralization of organic nitrogen to ammonium, which is preferentially used as a nitrogen source by microbes.

### Alphaproteobacteria (Rhizobiales) dominate peatland nitrogenase sequences

Diazotrophic methanotrophs have the potential to serve as a methane biofilter and a nitrogen source for peatland ecosystems. Previous work showed that *Alphaproteobacteria (Rhizobiales*), including Type II methanotrophs (63, 99), were the dominant diazotrophs in *Sphagnum-*dominated peatlands and may provide the unaccounted nitrogen input resulting from an imbalance in atmospheric nitrogen deposition and accumulation in *Sphagnum* mosses (44, 54, 60, 83, 99). However, the carbon metabolism of peatland diazotrophs remains unclear because the two dominant *Rhizobiales* families grow on both complex organics (*Bradyrhizobiaceae* (32)) as well as simple alkanes and C1 compounds (*Beijerinckiaceae* (19)). Consistent with previous studies, *Rhizobiales* showed the highest relative abundance in transcript libraries in this study. *Beijerinckiaceae* and *Bradyrhizobiaceae* dominated *nifH* and *nifD* amplicons, respectively. However, the complex taxonomic classification of *Rhizobiales nifH* and *nifD* genes (22) prevented distinguishing methanotrophic *Beijerinckiaceae* from heterotrophic *Beijerinckia indica* and *Bradyrhizobiaceae* solely on the basis of *nifH* and *nifD* phylogenies.

### Diazotrophy is catalyzed by the molybdenum form of nitrogenase

Peatland conditions such as low pH, nitrogen, and temperature would be expected to favor diazotrophy by alternative nitrogenases. Molybdenum sorption to peat is enhanced at low pH (6), biological requirements for Mo are higher when bacteria are fixing N_2_ than growing on other nitrogen compounds (33), and alternative nitrogenases have higher activity and expression at lower temperatures (74, 101). Below 10 nM Mo, diazotrophy is limited in laboratory cultures (1, 26, 34, 87, 108) and alternative nitrogenases, if present, are expressed (3). Transcription of alternative nitrogenase genes has been reported for *Peltigera* cyanolichens (45), but to our knowledge has not previously been investigated for *Sphagnum* peatlands.

Molybdenum concentrations in S1 bog porewaters were 1-7 nM, which is within the same range as Mo in other oligotrophic freshwaters (33), and lower than V (5-21 nM) and Fe (7-35 μM). Similar metal concentrations between: (a) sampling dates in September 2014, June 2015, and September 2015; (b) in another ombrotrophic Minnesota peatland, Zim bog, ∽80 km from the S1 bog; and (c) in acidic peatlands in Northern Europe (53) suggest that the values we measured are spatially and temporally representative for diverse northern peatlands.

Intriguingly, despite the presence of conditions that would be expected to favor alternative nitrogenase expression (low pH, long winters, and Fe>V>Mo), our collective evidence suggests that diazotrophy at the S1 bog was catalyzed by the Mo-containing form of the nitrogenase enzyme. The majority of *nifH* genes retrieved from metagenomes belonged to Mo-containing clusters I and III. A significant number of sequences came from uncharacterized Cluster IV, recently shown to contain a functional nitrogenase (110) that likely binds a Mo-Fe cofactor (72). The metagenomes contained very few cluster II *vnfD/anfD* genes, and minimal *vnfD / anfD* transcripts were amplified from peat cDNA. Ethane, a biomarker of alternative nitrogenase, was undetectable in ARA incubations. Finally, the ^15^N_2_-to-ARA conversion factor (3.9) was within the same range (3-4) as other peat bogs (41, 62, 99), and matched Mo-nitrogenase in pure culture experiments as opposed to the lower values measured for alternative nitrogenases (4). The question of how peatland diazotrophs access scarce Mo remains uncertain; it is possible that Mo bound to peat organic matter can be scavenged by diazotrophs as is the case in forest soils (71, 104).

### Methanotrophs are active in incubations and inhibited by acetylene

We performed bottle experiments to test whether methanotrophs were active. Complete CH_4_ consumption in air amended with 1% CH_4_ showed that methanotrophs were active in our incubations, and that the black bromobutyl stoppers we used were not toxic to peatland methanotrophs, whereas non-halogenated stoppers are toxic to aquatic aerobic methanotrophs (76). Acetylene fully inhibited CH_4_ consumption, demonstrating that the methanotrophs in our incubations were C_2_H_2_ sensitive, similar to laboratory strains tested in previous studies (16, 94). Like diazotrophy, methanotrophy was apparently also mediated by the enzyme requiring the scarcer metal; dissolved Fe was consistently orders of magnitude higher than Cu in peat porewater, yet *pmoA* sequences were more abundant than *mmoX* sequences in surface peat where the highest CH_4_ consumption was observed (24, 68). This finding is also consistent with higher transcription of *pmoA* vs. *mmoX* in other acidic peatlands (10, 53, 63). In laboratory studies, the “copper switch” from sMMO to pMMO supported growth occurs when Cu > 1 μM (90), which is several orders of magnitude higher than Cu concentrations measured at the S1 bog (7-38 nM). It is possible that the “copper switch” occurs at a lower threshold in peat bogs, or that there are other factors controlling the type of MMO expressed, such as CH_4_ or O_2_ availability. Additionally, the inherent nature of the peat matrix, with characteristically high levels of particulate and dissolved organic matter, likely also affects metal bioavailability in complex ways (98) not addressed in our study.

### Surface peatland diazotrophy is sensitive to oxygen and acetylene

If methanotrophs were the dominant diazotrophs in peatlands, as previously proposed (60, 99), CH_4_ addition should have stimulated ^15^N_2_ incorporation in our bottle experiments, and C_2_H_2_ should have inhibited it. Instead, this and previous work (61) found that ^15^N_2_ incorporation was not affected by CH_4_ addition. Acetylene partially inhibited ^15^N_2_ incorporation under degassed conditions (as in Kox, et al. (59)) and had minimal effect on oxic ^15^N_2_ incorporation, suggesting that C_2_H_2_-and O_2_-sensitive microbial clades contributed approximately half of the diazotrophic activity in our incubations. This finding highlights the importance of quantifying peatland diazotrophy by ^15^N_2_ incorporation instead of, or in addition to, ARA.

Based on *nif, pmoA* and *mcrA* phylogeny from native peat, the O_2_-and C_2_H_2_-sensitive diazotrophs were likely methanotrophic *Beijerinckiaceae* that can only fix N_2_ under micro-oxic conditions, and/or strict anaerobes in cluster III, such as methanogenic *Euryarchaeota*. These two families are the most active in surface peat based on transcript and amplicon sequences. Since *nifH* transcripts were 1-4 orders of magnitude lower in our incubations than native peat, it is likely that diazotrophs were stressed, possibly due to O_2_ exposure during the sampling process. Indeed, we observed highest ARA rates in hollows where the water table was typically at the bog surface, limiting O_2_ penetration into the peat. The other half of O_2_-sensitive diazotrophic activity was likely performed by C_2_H_2_-insensitive, heterotrophic *Bradyrhizobiaceae* and/or *Beijerinckiaceae* (51, 91), or C_2_H_2_-insensitive, methanotrophic *Methylocystaceae*, which can adapt to a wide range of CH_4_ concentrations (17). Less O_2_-sensitive microbes, including heterotrophic *Beijerinckiaceae* (91) and/or photosynthetic *Oscillatoriophycideae* (25), likely contributed to the minor amount of ARA activity in the presence of O_2_.

### Molecular markers for diazotrophy

We end with a word of caution with regard to the molecular detection of diazotrophs. The majority of studies in peatlands have employed PCR amplification and sequencing of the *nifH* marker gene for studying the dynamics of diazotrophs in peatlands. A wide range of *nifH* primer sets exist, with varying universality (29). Peat bog sequencing efforts have used polF/polR (this study; 63), F1/R6 (99), FGPH19/polR + polF/AQER (61), and 19F/nifH3 + nifH1(1)/nifH2(2) with nested PCR (8, 59). *In silico* evaluation predicts that the polF/polR primer set will not amplify the majority of *Proteobacteria* and/or *Cyanobacteria* and Group III *nifH* sequences (29), however this primer set yields the highest efficiency for qPCR (30). Of the *nifH* primer sets used previously, F1/R6, 19F/nifH3 and nifH1(1)/nifH2 are predicted to have the highest coverage for soils (>80% predicted primer binding for sequences from soil ecosystems). However, it is important to be aware that the F1/R6 primer set contains a number of mismatches with cluster III in peatlands, including methanogenic *Euryarchaeota* represented (Fig. S7). In order to maximize sequence coverage, we suggest primer sets that can amplify *nifH* from cluster III, such as IGK(3)/DVV (29), for future studies.

### Conclusions

This study revealed that peatland diazotrophs preferentially transcribed the Mo-based, rather than the V-based, form of the nitrogenase enzyme, despite the dominance of V over Mo in the environment. It also highlighted the sensitivity of diazotrophic peatland communities to O_2_ exposure during sample collection, and quantified the inhibitory effect of C_2_H_2_ addition on peatland diazotrophy. Under our experimental conditions in lab incubations, we did not observe CH_4_-stimulated diazotrophy. However, quantification of the relative contributions of methanotrophic and heterotrophic diazotrophy *in situ* awaits further investigation.

## Acknowledgments

This research was funded by DOE support to J.E.K. under grant numbers DE-SC0007144 and DE-SC0012088, and was supported by NASA Exobiology grant NNX14AJ87G to J.B.G. We thank Loren Dean Williams for inspiration with visuals, Heather Dang (UC Berkeley) for IRMS analysis, and Will Overholt for technical assistance. Work at LLNL was conducted under the auspices of DOE Contract DE-AC52-07NA27344, with funding provided by LDRD 14-ER-038.

